# Spike-wave discharges during low-current thalamic deep brain stimulation in mice

**DOI:** 10.1101/2024.05.10.593558

**Authors:** Francisco J. Flores, Isabella Dalla Betta, John Tauber, David Schreier, Emily P. Stephen, Matthew A. Wilson, Emery N. Brown

## Abstract

**Background:** Deep brain stimulation of central thalamus (CT-DBS) has potential for modulating states of consciousness, but it can also trigger spike-wave discharges (SWDs).

**Objectives:** To report the probability of inducing SWDs during CT-DBS in awake mice.

**Methods:** Mice were implanted with electrodes to deliver unilateral and bilateral CT-DBS at different frequencies while recording EEG. We titrated stimulation current by gradually increasing it at each frequency until an SWD appeared. Subsequent stimulations to test arousal modulation were performed at the current one step below the current that caused an SWD during titration.

**Results:** In 2.21% of the test stimulations (10 out of 12 mice), CT-DBS caused SWDs at currents lower than the titrated current, at currents as low as 20 μA.

**Conclusion:** Our study found a small but significant probability of inducing SWDs even after titration and at relatively low currents. EEG should be closely monitored for SWDs when performing CT-DBS in both research and clinical settings.

## Introduction

Deep brain stimulation of central thalamic nuclei (CT-DBS) is an effective technique for regulating brain arousal: it has helped to speed recovery from minimally conscious states (MCS) in humans [1, 2], reverse the effects of anesthesia in non-human primates (NHPs) [3–5], and enhance cognitive abilities in awake rodents and NHPs [6–9]. The neurons in the central thalamus are uniquely positioned to modulate brain arousal states as they send excitatory projections to the striatum and frontal and association cortices, and traumatic brain injuries involving central thalamic nuclei result in a variety of disorders of consciousness, including MCS [10].

Electrical stimulation of the central thalamus can also induce spike-wave discharges (SWDs) in cats [11], NHPs [12, 13], and humans [14]. In NHPs, CT-DBS has also been reported to induce a “vacant stare” that resembles absence seizures [15]. SWDs are characterized by a series of spikes and waves observed in the electroencephalogram (EEG) or local field potentials and are associated with absence seizures in humans [16]. SWDs are not only produced by central thalamic electrical stimulation but also by optogenetic stimulation of this area [17]. In this study we report the probability of inducing SWDs during test stimulations performed at the current one step below the current that caused an SWD during titration.

## Methods

### Animals and Surgery

All experiments were approved by the MIT IACUC (protocol # 2303000480). C57BL/6J mice were implanted (n = 12, 7 male) under isoflurane anesthesia with bipolar tungsten electrodes for stimulation (∅ = 125 μm, P1) aimed at the central thalamus (AP: −1.0 to −1.4, ML: *±*0.7, DV: −2.6 to −2.7; implanted impedances ≤200 kΩ), and stainless steel screws for EEG recording (8209, Pinnacle) were implanted over the frontal (AP: +1.5, ML: +0.7) and parietal (AP: −3.0, ML: −1.3) cortices, with ground and reference screws over the cerebellum.

### Recording and Stimulation

The EEG was recorded at 2713 Hz and band-passed (0.5–500 Hz; Neuralynx). Constant current stimulation (STG-8000, Multichannel Systems) consisted of biphasic, cathodal first, charge-balanced, 100 μs square pulses separated by a 100 μs isoelectric period and were applied either unilaterally or bilaterally [18].

### Experiments

Titration consisted of 10-s stimulation periods separated by 10-s off periods at gradually increasing currents until an SWD was observed or 200 μA was reached (Fig. 1A, left). 60-s test stimulations, separated by 120-s off periods, were performed at the current one step below the current that caused an SWD (Fig. 1A, right). Titration and test stimulations were repeated at each of five frequencies (1, 10, 50, 100, and 200 Hz).

**Figure 1:**
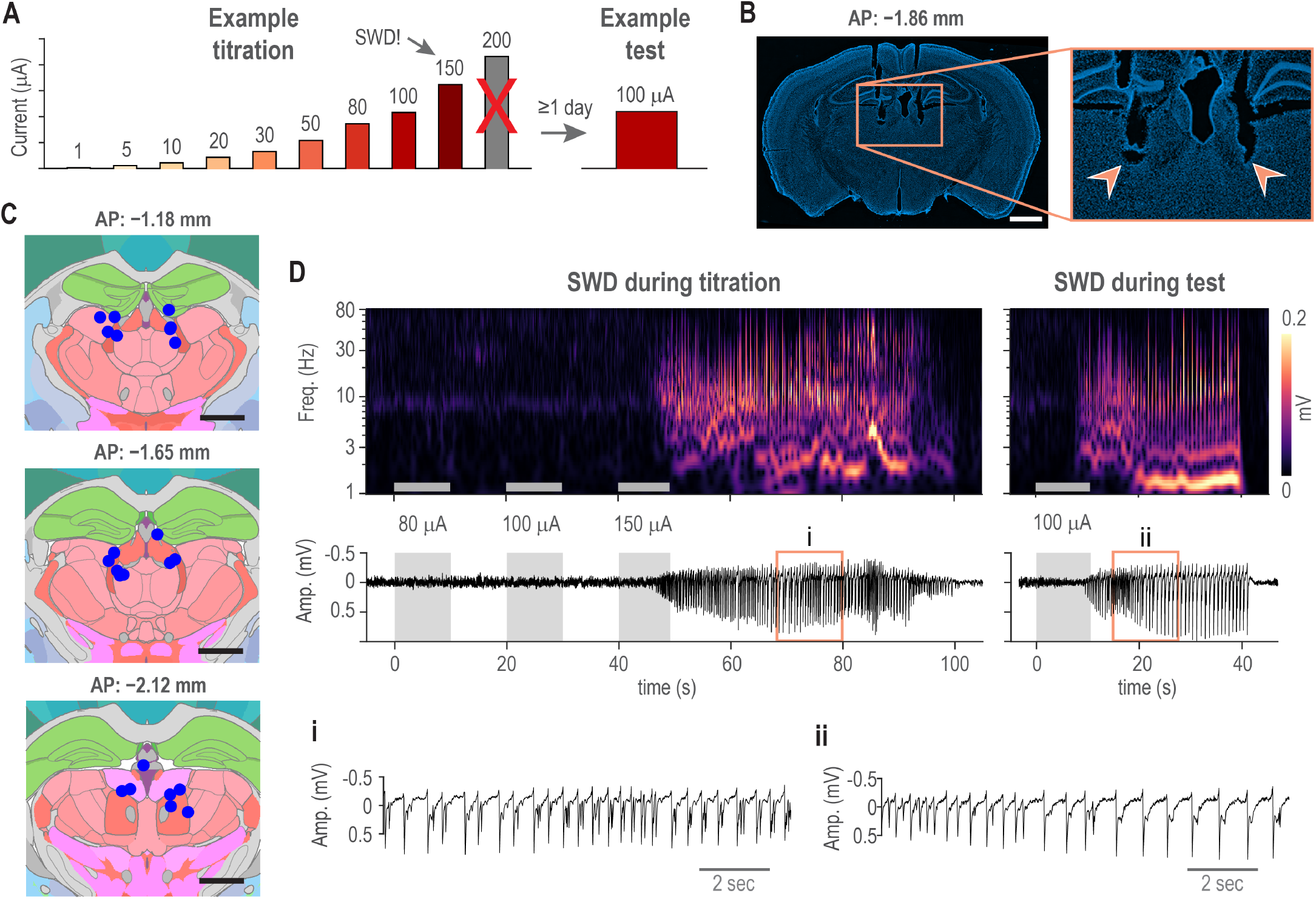
Methods and example SWDs induced by CT-DBS. A) Schematic of an example titration-test protocol at a given frequency. During titrations, 10-s stimulations at the currents shown were performed until an SWD was observed or 200 μA was reached. For example, if an SWD occurred at 150 μA, the titration was halted, and the subsequent 60-s test was performed at 100 μA. B) Representative example of a DAPI-stained coronal slice at AP −1.86 mm from bregma with zoomed-in view of the electrolytic lesions, indicated with arrowheads. C) Summary of all electrode tip locations (blue dots) projected onto the closest of three coronal planes, AP −1.18, −1.65, and −2.12 mm. D) Example of a titration-test pair at 200 Hz. The scalogram (top) and parietal EEG (bottom) show an SWD beginning during titration stimulation at 150 μA (left panel) and an SWD beginning during the subsequent test stimulation at 100 μA (right panel). Zoom-in of the SWDs observed in the parietal EEG (orange box) during titration (i) and during test (ii).

### Histology

After experiment completion, electrolytic lesions were created using 20 μA DC current for 10 s in anesthetized mice. Later, the mice were anesthetized and perfused with 10% formalin. The brains were sliced in 50-μm sections, mounted using DAPI-containing media, and imaged using a fluorescence microscope and the Zen Blue software (Zeiss). To locate the electrode tip, coregistration to the Allen Mouse Brain CCF [19] was performed in MATLAB using open-source software [20].

### Data Analysis

Scalograms were computed using the continuous wavelet transform (CWT) with an analytic morlet wavelet as implemented in MATLAB. All EEG recordings were independently reviewed by two researchers and SWDs present in at least one electrode were identified. Discrepancies were resolved by a third researcher. Summary statistics were computed as median and median absolute deviation (MAD). Analysis of SWD features was performed using a generalized linear mixed-effects model [21] as implemented in the lme4 R package [22], with SWD duration, latency, or probability as responses; mice as random intercepts; and sex, laterality, stimulation frequency, and stimulation current as predictors. A normal distribution was used to model duration, gamma to model latency, and binomial to estimate probability. In all models, frequency and current were log_10_-transformed. The effect of previous SWDs (within-day and across-day) was assessed but were excluded from final models based on a process of model selection using Akaike Information Criteria (AIC) [23].

Because the rate of SWD occurrence was near zero, confidence intervals for empirical probabilities were calculated using the Wilson score [24] rather than the typical Wald method.

## Results

We implanted the stimulation electrodes aiming to cover the central thalamus along the anteroposterior axis. A representative DAPI-stained slice shows the tracks and electrolytic lesions of an electrode pair (Fig. 1B). The AP coordinates of the electrode tips ranged from −1.0 to −2.4, covering several central thalamic nuclei (Fig. 1C, blue circles). Representative examples of SWDs occurring in a titration-test pair are shown in figure 1D. In this example, the SWD appeared during titration at 150 μA (Fig. 1D, left) and then during test at 100 μA (Fig. 1D, right). It is unlikely our protocol leads to spontaneous SWDs because out of 82 SWDs, only one did not begin within 15 s after stimulation onset. This SWD occurred 12.60 s after the offset of a previous SWD and was excluded from further analysis.

Considering that titrations were performed to find the highest current that would not cause an SWD for each mouse, we anticipated that test stimulations performed at lower currents would not cause SWDs. However, we found that 2.21% of test stimulations induced SWDs (Fig. 2A), at currents between 20 and 200 μA. SWDs were observed during test stimulations in 10 out of 12 mice (Fig. 2B, blue). However, estimation of individual probabilities is significantly higher than zero across the mice population (Fig. 2B, purple), which results in an overall significant probability of causing an SWD during a test stimulation (2.21%, CI [1.46, 3.33]) (Fig. 2C, dashed line). Higher currents during tests had a higher probability of inducing an SWD (*P* = 1.97 *·* 10^−4^, Fig. 2C).

**Figure 2:**
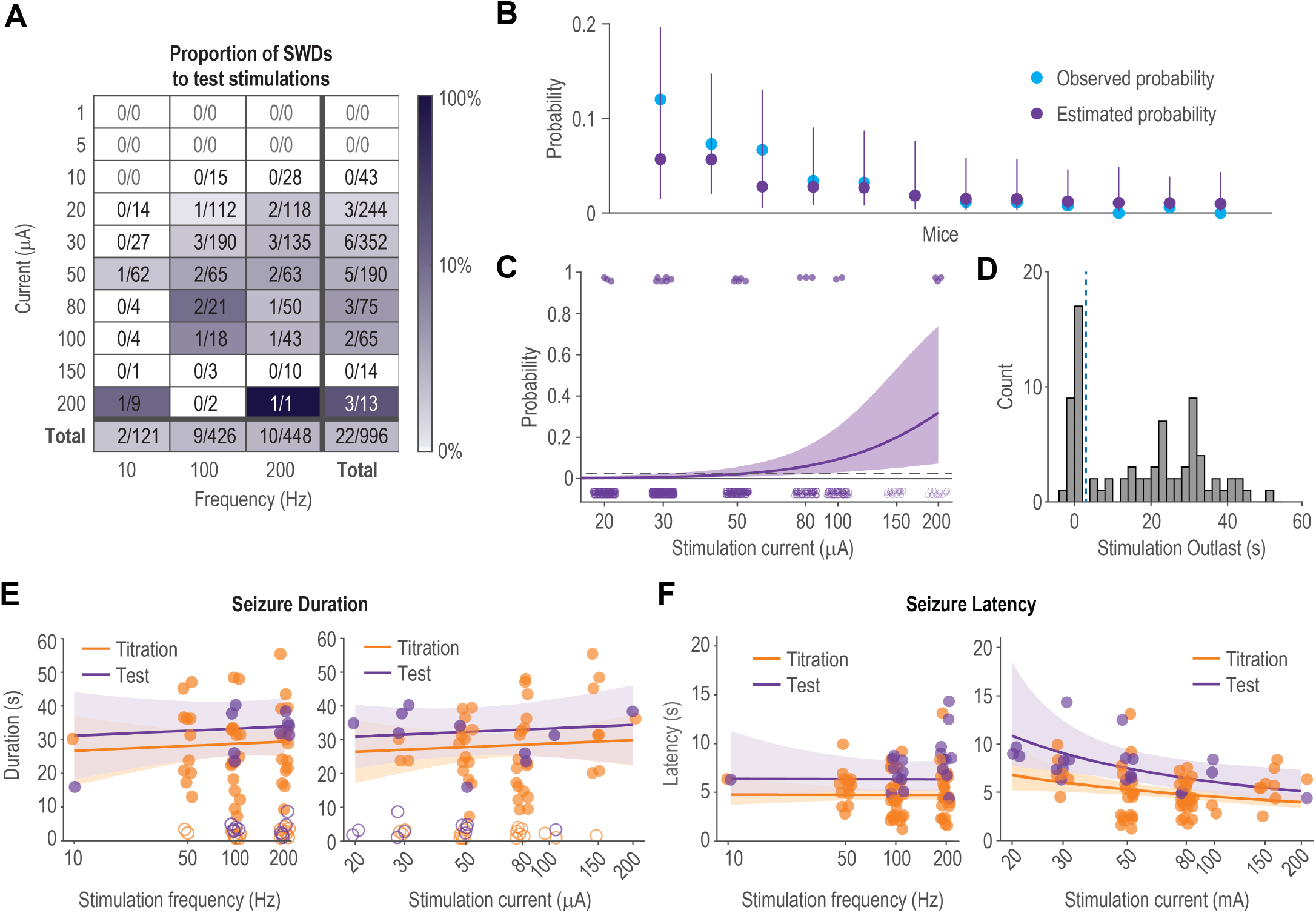
Probability and features of SWDs. A) Frequency by current matrix with the number of SWDs induced over the total number of stimulations performed. The percent of test stimulations that resulted in SWDs is indicated by the color scale. Totals for each current are shown in the final column, and totals for each frequency in the final row. The bottom right cell shows the grand total SWDs over the grand total of stimulations (2.21%). One stimulation (50 Hz at 200 μA) was excluded from the figure for simplicity but is included in the totals. B) Observed probability (blue dots), estimated probability (purple dots), and 95% CI (purple lines) of SWD during test stimulations for each mouse. C) Total probability (purple line) and 95% CI (purple shade) of an SWD occurring during test stimulations at different currents across all frequencies. The dots represent occurrence (upper) and absence (lower) of SWDs across all test stimulations. Dashed line represents total probability of SWD occurrence (2.21%). D) Histogram of the time each SWD outlasted the stimulation. The dashed line shows the 2 s cutoff. E) The relationship between frequency (left) and current (right) and SWD duration across titrations (orange) and tests (purple). F) Same as E but for SWD latency.

We compared SWD duration and latency to assess differences between SWDs during both titration and test stimulations. Very short SWDs (33%) which outlast stimulation by less than 2 second were excluded from further duration analysis (duration = 2.39 s, MAD = 0.99 s; Fig. 2D and Fig. 2E, open circles). The remaining 67% of SWDs outlasted the stimulation by more than 5 s (duration = 30.21 s, MAD = 7.41 s). Regarding SWD duration, there was no significant relationship with stimulation frequency (*P* = 0.66, Fig. 2E, left), current (*P* = 0.65, Fig. 2E, right), laterality of stimulation (*P* = 0.60), sex (*P* = 0.16), nor whether it was a titration or a test (*P* = 0.26, Fig. 2E). Median latency from stimulation onset to SWD onset was 5.63 s (MAD = 1.61 s). Stimulation frequency did not have an effect on SWD latency (*P* = 0.97, Fig. 2F, left), but current had a significant effect on SWD latency (*P* = 0.011, Fig. 2F, right), with shorter latencies at higher currents. There was also a significant difference in SWD latency between test and titration (*P* = 0.002, Fig. 2F), with shorter latencies associated with titrations. Sex and laterality of stimulation had no significant relationship to latency (*P* = 0.19, *P* = 0.25).

## Discussion

We have shown that there is a small but significant probability that CT-DBS can induce SWDs at stimulation currents as low as 20 μA. This occurred in 10 out of 12 mice, even after accounting for individual variability of SWD-inducing currents through titration. The probability of inducing an SWD during test stimulations increased with increasing current. We did not observe a significant difference in duration between SWDs induced by titration and test stimulations nor was there a relationship with stimulation current or frequency, likely because seizure stopping mechanisms are due to intrinsic homeostatic feedback [25] and are independent from stimulation parameters. SWD latency was significantly shorter for SWDs induced during titrations, which might be due to the shorter off periods between titration stimulations leading to increased concentration of extracellular potassium, which once elevated can take minutes to return to baseline [26]. Latency was also significantly shorter at higher currents, possibly due to a larger volume of effective stimulation recruiting more neurons [27], leading to faster SWD onset. We observed similar EEG characteristics in SWDs induced by titration and test stimulations, probably due to intrinsic properties of thalamocortical circuits [28].

SWDs are not reported to occur during CT-DBS in humans in MCS [1, 2] nor anesthetized NHPs [3–5], possibly because lesions causing MCS result in low levels of neural activity and most anesthetics suppress seizure activity. During wakefulness, CT-DBS causes SWDs in cats [11] and humans [14], and in NHPs it causes SWDs [12, 13] or behavior that resembles absence seizures [15].

Awake CT-DBS causes SWDs at different frequencies, currents (or voltages), pulse durations, stimulation durations, and lateralities. We tried both unilateral and bilateral stimulation at frequencies and a pulse duration consistent with previous reports [4], but we selected lower currents and shorter stimulation duration to diminish the chances of producing SWDs or tissue damage [18]. DBS may have both local and distant effects through stimulation of fibers passing near the electrode [27]. However, optogenetic stimulation of the central thalamus also produces SWDs and convulsive seizures in rats [17], suggesting that CT-DBS-induced SWDs are due to local thalamic stimulation rather than distant effects.

Our aim was to test modulation of brain states with CT-DBS, while avoiding SWDs by using currents lower than the SWD-inducing current determined during titration. However, we found there was experiment-to-experiment variability in SWD-inducing currents within mice which led to SWDs during test stimulations. This variability may be explained because the effects of DBS depend on the ongoing neural activity immediately preceding the stimulation [29]. Kindling effects are unlikely as we found that the history of previous stimulation, both within and across days, did not affect the probability of inducing an SWD. Considering our results, EEG should be closely monitored for SWDs when performing CT-DBS, especially in awake subjects.

## CRediT author statement

**Francisco J. Flores:** Conceptualization, Formal analysis, Investigation, Data curation, Writing – original draft, Writing – review & editing, Supervision. **Isabella Dalla Betta:** Conceptualization, Formal analysis, Investigation, Data curation, Writing – original draft, Writing – review & editing. **John Tauber:** Formal analysis, Writing – original draft, Writing – review & editing. **David Schreier:** Formal analysis, Writing – review & editing. **Emily P. Stephen:** Formal analysis, Writing – review & editing, Supervision. **Matthew A. Wilson:** Writing – review & editing, Supervision, Funding acquisition. **Emery N. Brown:** Conceptualization, Writing – review & editing, Supervision, Funding acquisition.

## Declaration of competing interest

E.N.B holds patents on anesthetic state monitoring and control. E.N.B. holds founding interest in PASCALL, a start-up developing physiological monitoring systems; receives royalties from intellectual property through Massachusetts General Hospital licensed to Masimo. The interests of E.N.B. were reviewed and are managed by Massachusetts General Hospital and Mass General Brigham in accordance with their conflict of interest policies.

## Acknowledgements

This work was generously supported by the JPB Foundation; the Picower Institute for Learning and Memory; George J. Elbaum (MIT ‘59, SM ‘63, PhD ‘67), Mimi Jensen, Diane B. Greene (MIT, SM ‘78), Mendel Rosenblum, Bill Swanson, annual donors to the Anesthesia Initiative Fund; and the NIH Awards P01 GM118269 and R01 NS123120 (to E.N.B.). D.S. is supported by the Swiss National Science Foundation (P500PM 210834) and the Swiss Neurological Society.

## Notes

### Competing Interest Statement

Emery N. Brown holds patents on anesthetic state monitoring and control. E.N.B. holds founding interest in PASCALL, a start-up developing physiological monitoring systems; receives royalties from intellectual property through Massachusetts General Hospital licensed to Masimo. The interests of E.N.B. were reviewed and are managed by Massachusetts General Hospital and Mass General Brigham in accordance with their conflict of interest policies.

